# Shared Genetic Architecture between Asthma and Allergic Diseases: A Genome-Wide Cross Trait Analysis of 112,000 Individuals from UK Biobank

**DOI:** 10.1101/133322

**Authors:** Zhaozhong Zhu, Phil H. Lee, Mark D. Chaffin, Wonil Chung, Po-Ru Loh, Quan Lu, David C. Christiani, Liming Liang

**Affiliations:** Department of Epidemiology, Harvard T.H. Chan School of Public Health, Boston, Massachusetts, USA.; Department of Environmental Health, Harvard T.H. Chan School of Public Health, Boston, Massachusetts, USA.; Center for Human Genetics Research, Massachusetts General Hospital and Harvard Medical School, Boston, Massachusetts, USA.; Psychiatric and Neurodevelopmental Genetics Unit, Massachusetts General Hospital and Harvard Medical School, Boston, Massachusetts, USA.; Medical and Population Genetics Program, Broad Institute of MIT and Harvard, Cambridge, Massachusetts, USA.; Stanley Center for Psychiatric Research, Broad Institute of MIT and Harvard, Cambridge, Massachusetts, USA.; Department of Biostatistics, Harvard T.H. Chan School of Public Health, Boston, Massachusetts, USA.

## Abstract

Clinical and epidemiological data suggest that asthma and allergic diseases are associated. And may share a common genetic etiology. We analyzed genome-wide single-nucleotide polymorphism (SNP) data for asthma and allergic diseases in 35,783 cases and 76,768 controls of European ancestry from the UK Biobank. Two publicly available independent genome wide association studies (GWAS) were used for replication. We have found a strong genome-wide genetic correlation between asthma and allergic diseases (*r_g_* = 0.75, *P* = 6.84×10^−62^). Cross trait analysis identified 38 genome-wide significant loci, including novel loci such as D2HGDH and GAL2ST2. Computational analysis showed that shared genetic loci are enriched in immune/inflammatory systems and tissues with epithelium cells. Our work identifies common genetic architectures shared between asthma and allergy and will help to advance our understanding of the molecular mechanisms underlying co-morbid asthma and allergic diseases.

## Introduction

Asthma is a chronic respiratory syndrome that is characterized by abnormal and inflamed mucosa of the airways, wheezing, and shortness of breath. Allergic diseases are immune responses for allergies, such as allergic rhinitis and atopic dermatitis (eczema). Asthma, allergic rhinitis and eczema all belong to type I hypersensitivity, which is an immune response to foreign antigen and often associates with immunoglobulin E (IgE)-mediated inflammation^1,2^. Genetic studies offer a structured means of understanding the causes of asthma and allergic diseases, as well as identifying targets that can be used to treat the syndrome^3-7^.

Clinical and epidemiological studies suggest that asthma and allergy are associated^8,9^. Several studies have identified allergic diseases, such as allergic rhinitis and eczema, as a risk factor for asthma, with the prevalence of allergic rhinitis in asthmatic patients being 80% to 90%^8,10^. These studies demonstrate that the coexistence of asthma and allergy is frequent, and that allergy usually precedes asthma. Also our previous epigenetic study has identified methylation loci linked to asthma and allergy via IgE pathway^11^.

One hypothesis to account for the similar symptoms and conditions is that these diseases share a common genetic etiology. Cotsapas et al discovered nearly half of loci in genome-wide association studies (GWAS) of an individual disease influence risk to at least two diseases, indicating the shared genetic architecture of immune-mediated inflammatory and autoimmune diseases^12^. As each of the shared or similar risk factors has strong genetic influences on disease risk, the observed clustering of multiple risk factors could be due to an overlap in the causal genes and pathways^13-20^. In addition, grouping variants by the traits they influence should provide insight into the specific biological processes underlying comorbidity and disease risk.

Clinical and epidemiological studies have found asthma and allergic diseases can occur either in the same individual or in closely related family members^21-24^, suggesting potential pleiotropic effect. The heritability has been estimated at varying between 35% and 95% for asthma^21,25^ and 33% and 91% for allergic rhinitis, 71% and 84% for atopic dermatitis (eczema), 34% and 84% for serum IgE levels, which suggests that the genetic contribution to them can be significant^21^. While Hinds and colleagues investigated the shared genetic etiology for 38 allergic diseases including asthma^26^, understanding has been limited to only 16 genome-wide shared susceptibility loci which were based on self-reported phenotype.

In order to increase our knowledge of shared genetic determinants influencing doctor diagnosed allergic diseases and asthma, and to potentially discover novel loci, we investigated the genetic commonality among asthma and allergic diseases (hay fever/allergic rhinitis or eczema). We conducted a large scale GWAS analysis based on these traits to explore genetic correlations and shared genetic components among these diseases using data from the UK Biobank, which is the largest and most complete European Biobank available at present. Replications were done in two independent public available GWAS studies, the GABRIEL asthma GWAS study^4^ and the Early Genetics & Lifecourse Epidemiology (EAGLE) eczema consortium study^27^.

## Methods

### Study population and design

Study participants were from the UK Biobank study, described in detail elsewhere^28-30^. In brief, the UK Biobank is a prospective study of > 500,000 people living in the UK. All people in the National Health Service registry who were aged 40–69 years and living <25 miles from a study center were invited to participate from 2006–2010. In total, 503,325 participants were recruited from over 9.2 million mailed invitations. Self-reported baseline data were collected by questionnaire, and anthropometric assessments were performed. For the current analysis, individuals of non-white ethnicity were excluded to avoid confounding effects. All participants provided informed consent to the UK Biobank.

Overall study design was shown in Figure 1. To identify genetic variants that contribute to doctor-diagnosed asthma and allergic diseases and link them with other conditions, we performed GWAS using phenotype measures in UK Biobank participants (N = 152,249). We removed 39,698 samples that are non-European, related, withdrew from UK Biobank or are missing phenotype information. Thus, a total of 112,551 European descents with high-quality genotyping and complete phenotype/covariate data were used for these analyses, including 25,685 allergic diseases (hay fever/allergic rhinitis or eczema), 14,085 asthma and 76,768 controls for the analysis. There were 53,094 males and 59,457 females in the population.

**Figure 1.**
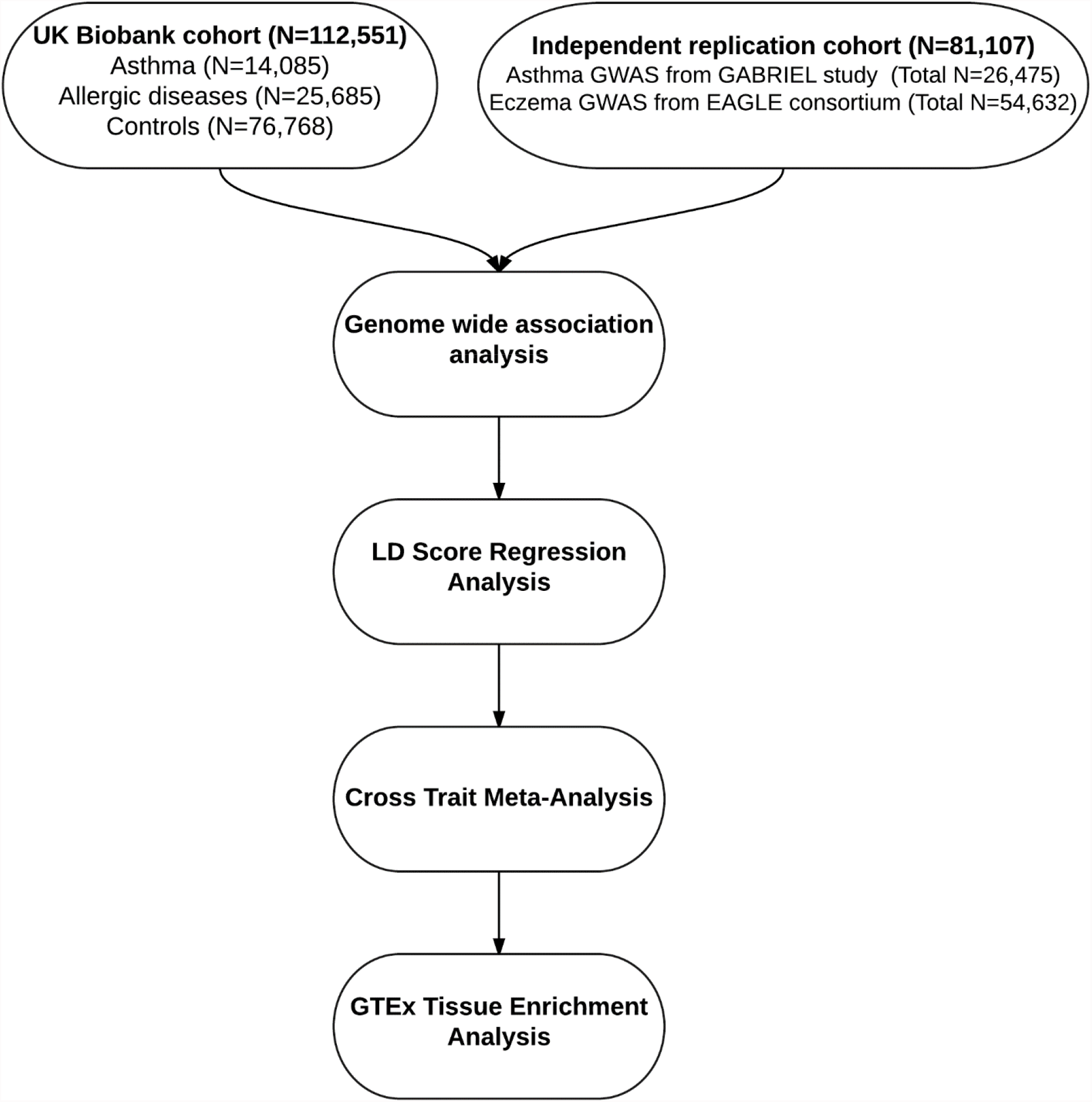
Overall study population and design. UK Biobank cohort consists of 112,551 subjects, including doctor diagnosed 14,085 asthma subjects and 25,685 allergic diseases, 76,768 controls. Two publicly available independent GWASs, an asthma GWAS (N = 26,475) and an eczema GWAS (N = 54,632) were used for replication.

Furthermore, we have included summary statistics for a replication analysis from an independent asthma GWAS (GABRIEL study, 10,365 cases 16,110 controls)^4^ and an independent eczema GWAS (EAGLE eczema consortium, 13,287 cases and 41,345 controls)^27^.

### Data summary, quality control (QC) and imputation

Detailed genotyping and QC procedures of UK Biobank were described previously (http://biobank.ctsu.ox.ac.uk/) and in supplementary material. In summary, after QC procedures were applied, the current released UK Biobank data contains 806,466 SNPs that passed SNP QC in at least one batch. Post-imputation QC was performed as previously outlined (http://biobank.ctsu.ox.ac.uk/); and genotype imputation procedure provided a total of 73,355,677 SNPs. In addition, we performed stringent QC standards by PLINK 1.90. ^31^ We selected variants that did not deviate from Hardy-Weinberg Equilibrium (HWE) (P>1×10^−12^), per variant missing call rates<5%, per-sample missing rate<40%, minor allele frequency (MAF)>1% and an imputation quality score (INFO) > 0.8. Quantile-Quantile (QQ) plots were produced and checked for each phenotype. R package GenABEL was used to compute genomic inflation values. A total of 7,489,529 SNPs passed QC on whole genome and with complete information in all three phenotypes, which were eligible for statistical association analyses.

### GWAS analysis

We performed additive logistic regression genetic association analysis adjusting for age, sex, genotyping array and ten ancestry principal components using PLINK 1.90^31^ to assess association between phenotype and genotype on each individual disease. After association analysis, we applied PLINK clumping function (parameters: –clump-p1 5e-8 –clump-p2 1e-5 – clump-r2 0.2 –clump-kb 500) to determine top loci that are independent to each other, i.e. variants with p-value less than 1×10^−5^, has r^2^ more than 0.2 and less than 500 kb away from the peak will be assigned to that peak’s clump. The top loci that were annotated to the same gene were then further combined as one independent loci for downstream analysis. We have used GWAS catalog (https://www.ebi.ac.uk/gwas, search date: 4/1/2017) to identify previously reported genes by other GWAS studies.

### LD score regression analysis

Post-GWAS genome-wide genetic correlation analysis by LD score regression (LDSC) ^32^ was conducted using all UK Biobank SNPs after imputation of 1000 Genomes Project variants and QC ^33^. LDSC estimates genetic correlation between two traits (ranging from −1 to 1) from summary statistics using the facts that the GWAS effect size estimate for each SNP incorporates the effects of all SNPs in linkage disequilibrium with that SNP. SNPs in a high linkage disequilibrium region would have higher χ2 statistics than SNPs in a low linkage disequilibrium region, and a similar relationship is observed when single-study test statistics are replaced with the product of the *z* scores from two studies of traits with some correlation ^32^. LDSC applied a self-estimated intercept during the analysis to account for shared subjects between studies^32^.

### Cross trait meta-analysis

After assessing genetic correlations among all traits, we applied cross trait GWAS meta-analysis by using the R package Association analysis based on SubSETs (ASSET) to combine the association evidence for asthma and allergic diseases at individual variants^34^. This method combines effect estimate and standard error of the GWAS summary statistics to test hypothesis of association between the SNP with any subset of studies. It can also account the correlation among studies/subjects that might arise due to shared subjects across distinct studies or due to correlation among related traits in the same study by using case-control overlap matrices.

### GTEx Tissue Specific Expression Analysis

In order to determine if shared asthma and allergy genes are overly expressed in disease-relevant tissues, we conducted tissue specific expression analysis (TSEA)^35,36^. Gene lists were generated from lists obtained from analyses for asthma, allergic diseases and cross-disease, and by including those which have a matching HUGO Gene Nomenclature Committee (HGNC) name. Genes identified in this way were analyzed for tissue enrichment using publicly available expression data from pilot phase of the Genotype-Tissue Expression project (GTEx)^37^ (www.gtexportal.org, 1/31/2013). In the GTEx project, postmortem samples from a wide variety of tissues and donors have been used for bulk RNA sequencing according to a unified protocol. All samples were sequenced using Illumina 76 base-pair paired-end reads. The pilot phase of the GTEx project consists of 1,839 samples derived from 189 post-mortem subjects, including samples from 45 different tissues, with some tissues offering multiple ‘sub-tissue’ types. TSEA averaged collapsed reads per kilobase per million mapped reads (RPKM) values for the sub-tissue types (25 ‘whole-tissue’ types). We filtered for unique HGNC IDs (n = 213 shared genes mapped by identified loci from meta-analysis result existing in tissue expression dataset). For each tissue, transcripts from the processed GTEx transcripts that are specifically expressed or enriched have been identified by using the TSEA pSI R package function to calculate the specificity index probability (pSI). Significance level of shared genes between asthma and allergic diseases enriched in each tissue were identified by by Fisher’s exact test with Benjamini-Hochberg correction.

### Replication analysis

In order to confirm our findings, we have further included two public available independent GWAS data: asthma GWAS, the GABRIEL consortium study ^4^ and eczema GWAS, the EArly Genetics & Lifecourse Epidemiology (EAGLE) eczema consortium study ^27^, for LD score regression analysis, cross trait analysis and GTEx tissue expression enrichment analysis. We have used LiftOver tool (http://genome.sph.umich.edu/wiki/LiftOver) to convert asthma GWAS reference genome from hg18 to hg19. ImpG-Summary was used to impute asthma GWAS summary statistics to the 1000 Genomes Project variants (phase 3 release v5)^38^. SNPs with imputation quality r2pred < 0.8 were removed. Cross trait analysis between two replication GWAS datasets was conducted using R package Cross-Phenotype Meta-Analysis (CPMA), which tests for the hypothesis that each independent SNP has multiple phenotype association and combines the empirical evidence based on their P-values^12^.

### Overrepresentation Enrichment Analysis

In order to understand the biological insights of the shared genes, we have used the WebGestalt tool ^39^ to assess overrepresented enrichment of the identified shared gene set between asthma and allergic diseases in KEGG^40^ pathways and Gene Ontology (GO) biological process^41,42^.

### Sensitivity analysis

To estimate the effect of using non-allergic asthma individuals, we performed additive logistic regression analysis adjusting for the same covariates as the primary analysis to assess the association between phenotype and genotype on each individual disease using only nonoverlapping samples between asthma and allergic diseases from the UK Biobank (excluding overlapped cases from asthma and allergic diseases, N = 6,177). We carried out cross-disease heritability analysis and meta-analysis to identify the loci that are associated with both diseases. Due to the large reduction in sample size of asthma cases and allergic diseases cases, the power of detecting association between phenotypes and SNP is expected to decrease.

### Cross tissue heritability of gene expression

The GTEx project^37^ (phs000424.v6.p1) was also used to quantify the cis SNP cross-tissue heritability 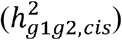 of gene expression across a range of tissues. All estimates were obtained using restricted maximum likelihood (REML) as implemented in Genome-wide Complex Trait Analysis (GCTA) 43. Only common variants that fall within 1 MB of the transcription start site of a gene were considered in deriving these estimates. Enrichment of 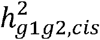 was determined by comparing average 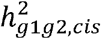 for a given gene list to 1,000,000 random gene lists of equal size in the same tissue pair.

## Results

The genetic correlation was calculated using common SNPs. There was strong positive genetic correlation between asthma and allergic diseases (*r_g_* = 0.75, *P* = 6.84×10^−62^). We have also extended our LDSC analysis to another type of immune complex mediated (Type III) disease: rheumatoid arthritis^44^, and two types of delayed cell mediated (Type IV) immune diseases: Crohn’s disease and ulcerative colitis^45^; however we did not find significant genetic correlation between asthma and other immune traits (rheumatoid arthritis, Crohn’s disease and ulcerative colitis), we confirmed the high genetic correlation between Crohn’s disease and ulcerative colitis (*r_g_* = 0.57, *P* = 7.98×10^−23^) (Table 1). Thus this empirical evidence of shared genetic etiology for asthma and allergic diseases encourages the investigation of common pathophysiology. Estimates of the SNP-based heritability by using GWAS summary statistics are presented in Supplementary Table 1. The proportion of variance in the traits explained by common genetic variants at the observed scale is 7.2% (s.e. 0.7%) for asthma and 7.5% (s.e. 0.7%) for allergic diseases.

**Table 1.**
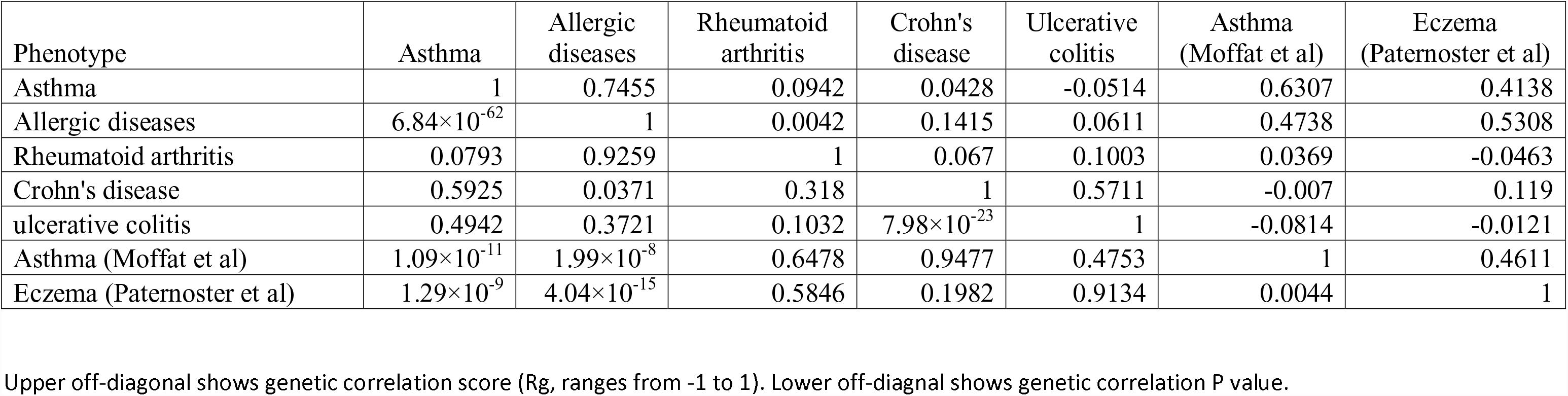
Genetic correlation between immune diseases.

The phenotype-genotype association test was carried out on all 112,551 samples and 7,489,529 SNPs from UK Biobank after QC. We identified 32 genome-wide significant (*P* < 5×10^−8^) independent loci for asthma and 33 for allergic diseases (See Method, Supplementary Table 2 and 3, Supplementary Fig. 4 and 5). The genomic control parameter λ was 1.13 for asthma and 1.14 for allergic diseases (Supplementary Fig. 1 and 2)^46-48^.

**Figure 2.**
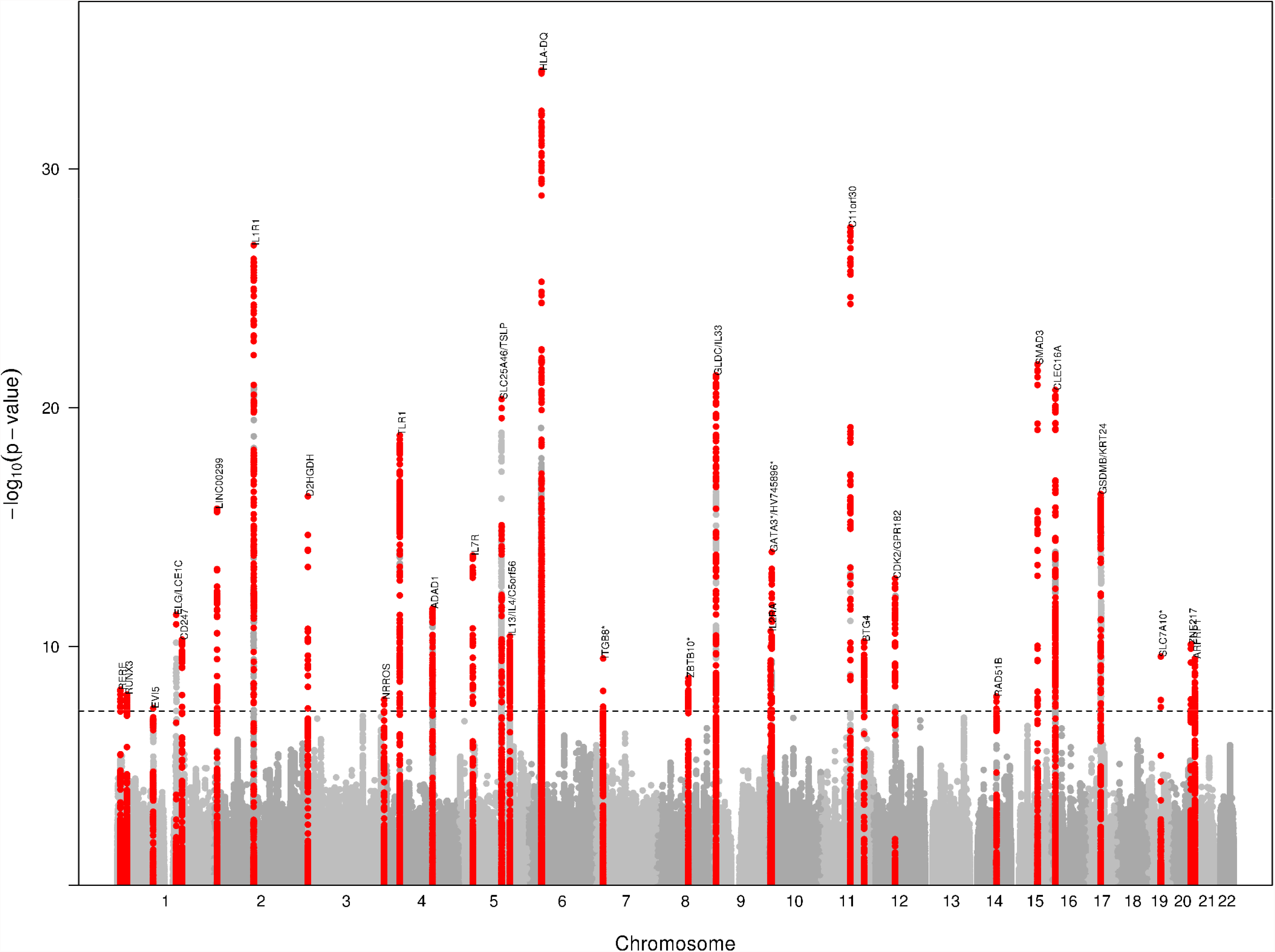
Manhattan plot of cross trait meta-analysis. The horizontal axis illustrates chromosome position, the vertical axis illustrates significance level −log_10_(P-value) of asthma and allergic diseases’ shared markers. Total of 38 independent loci was identified to be associated with both diseases. Dashed black line indicate genome-wide significance (*P* < 5×10^−8^)

For asthma, we confirmed most of the previously reported association loci, including IL1R1, IL1RL1, IL18R1, IL13, SLC25A46, HLA regions, SMAD3, GSDMB and many others (Supplementary Table 2). Out of the 32 independent loci reported here, 7 are novel associations with asthma and 4 of these new loci were within protein-coding genes body (Supplementary Table 2). For allergic diseases, genes such as C11orf30, FLG, SLC25A46, TMEM232, CAMK4, TSLP, WDR36, CLEC16A, DEXI, HLA regions and many others were confirmed in our analysis (Supplementary Table 2). Out of the 33 independent loci identified for allergic diseases, 11 are novel, of which 8 were mapped to protein-coding genes (Supplementary Table 3). Six SNPs were found to exactly overlap between asthma and allergic diseases (rs34290285 on D2HGDH and GAL3ST2, rs7705653 on SLC25A46 and TMEM232, rs12413578 near HV745896, rs7936070 on C11orf30, rs56062135 on SMAD3 and rs10414065 near SLC7A10), and 22 loci contains the same protein-coding genes between asthma and allergic diseases even though not at the same peak SNP.

Among 6 exactly overlapping SNPs, 4 were mapped to protein-coding genes. TMEM232 and SLC25A46 (rs7705653) encode transmembrane protein 232 that belongs to tetraspanin family and a solute carrier protein involved in transport of metabolites respectively. C11orf30 (rs7936070) is associated with total serum IgE levels in asthma^49^; also the locus C11orf30 increases susceptibility to poly-sensitization^50^. SMAD3 (rs56062135) is involved in the development of asthmatic inflammation due to responses of immune cells, such as cytokines and T-helper 2 (Th2) cells, which are known to be important in generating an inflammation that characterizes asthma and allergic disease^51^. The fourth SNP, rs34290285 falls on two novel genes for asthma and allergic diseases, D2HGDH and GAL3ST2, which are responsible for encoding metabolism related D-2hydroxyglutarate dehydrogenase enzyme^52^ and tumor metastasis related Galactose-3-O-sulfotransferase 2 enzyme^53^.

Furthermore, we have strengthened this finding in a cross-trait analysis and identified 38 loci contain SNPs with P < 5 ×10^−8^ (Fig. 2 and Table 2). The genomic control parameter λ is 1.31 for cross trait meta-analysis (Supplementary Fig. 3). Figure 2 shows the Manhattan plot of the cross-trait analysis between asthma and allergic diseases. The strongest association signal was observed on chromosome 6 at HLA-DQ region (rs9273374, *P*meta = 7.87×10^−35^)(Fig. 2 and Table 2), the first asthma-susceptibility locus identified^54^, and extended haplotypes encompassing HLA-DQ and HLA-DR have been studied for their effects on specific allergen sensitization^26,55^. The second strongest signal was mapped to C11orf30 on chromosome 11q13 (rs7936070, *P*_meta_ = 2.81×10^−28^) (Fig. 2 and Table 2), a gene associated with total serum IgE levels and increased susceptibility to poly-sensitization. This SNP was also an overlapping SNP between asthma and allergic diseases single GWAS model. The third strongest signal was observed on IL1R1 genes (rs72823641, *P*_meta_ = 1.58×10^−27^)^56^ (Fig. 2 and Table 2). Cytokine receptors have an impact on many cytokine induced immune and inflammatory responses, such as asthma and allergy^4,27,57^.

**Table 2.**
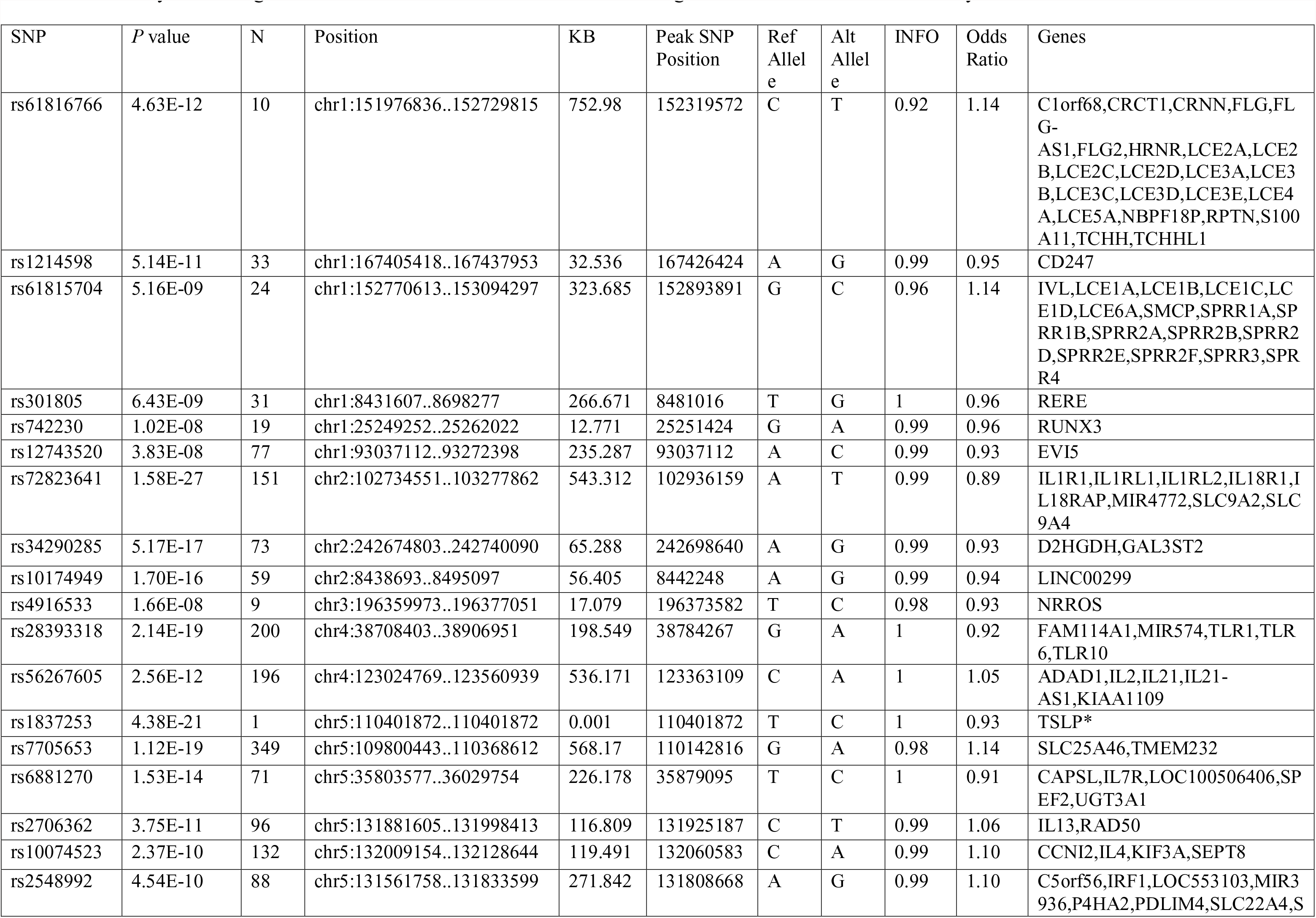
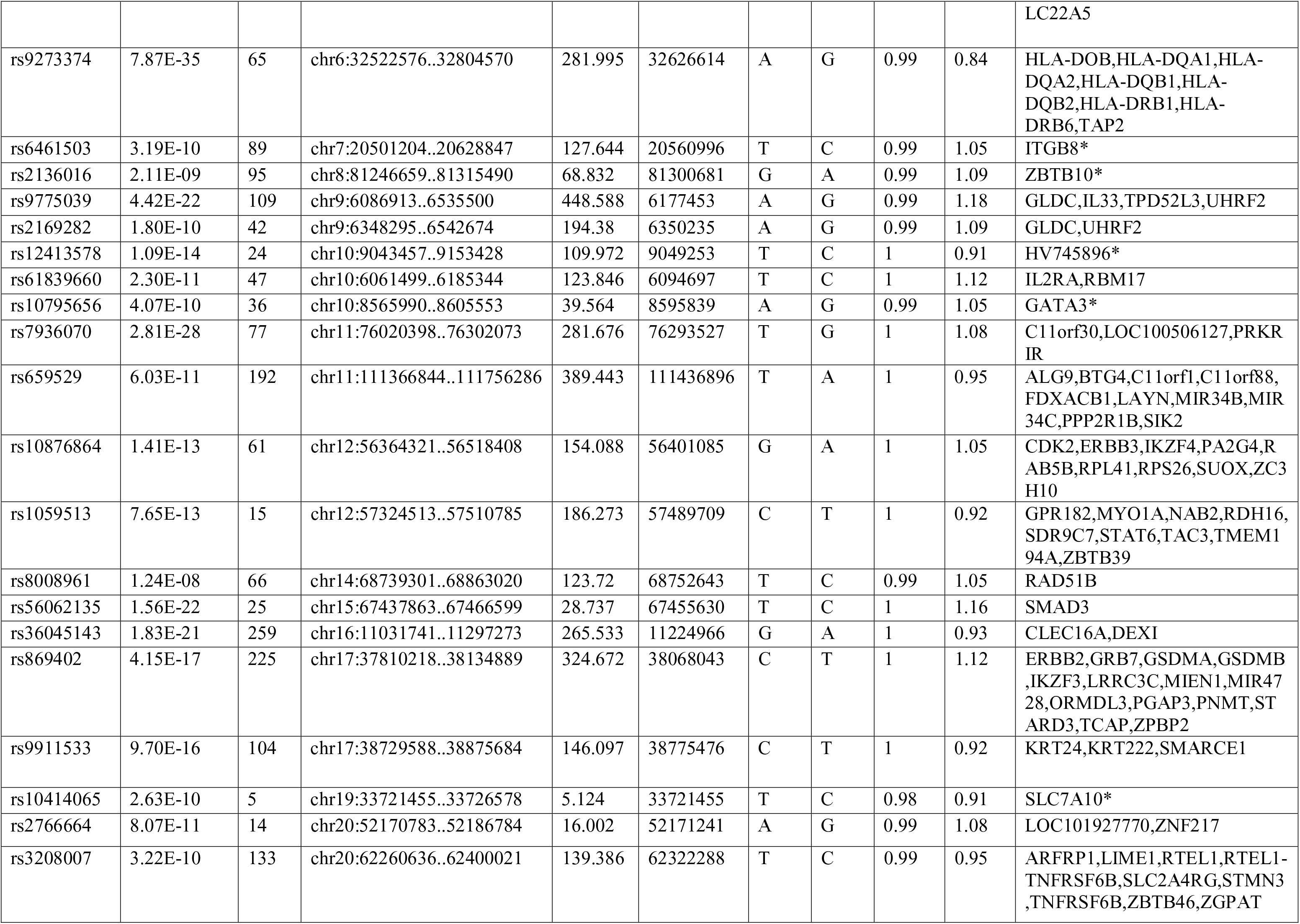
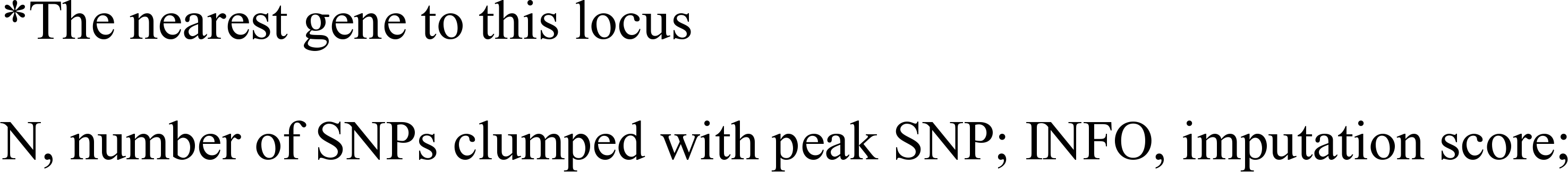
Summary of the 38 genomic loci associated with the asthma and allergic disease in cross trait meta-analysis.

The fourth strongest signal observed on SMAD3 (rs56062135, *P*_meta_ = 1.56×10^−22^) is another loci found in both asthma and allergic diseases, which is known to have a role in immune cell response^51^. We have also found the FLG (Filaggrin) gene (rs61816766, *P*_meta_ = 4.63×10^−12^) to be associated with both asthma and allergic diseases, which is crucial for maintaining normal skin layer function^58^. Our cross-trait results showed most loci are significant for both diseases, and suggests that it is these shared genetic loci between two diseases driving the overall significant positive genetic correlation.

To understand whether the 38 loci as a group are enriched for expression in certain tissue types, we used the GTEx pilot data^37^. The GTEx TSEA enrichment analysis identified five independent tissues expression that were significantly enriched (after Benjamin-Hochberg correction) for expression of cross-trait associated genes (Fig. 3). The most strongly enriched tissue was part of the integumentary system (skin). The other significantly enriched tissues included esophagus, vagina, lung and whole blood. We also found that the identified shared genes between asthma and allergic diseases have significant cross tissue heritability between each pair of tissues, particularly skin and lung have relatively high cross tissue heritability (Supplementary Fig. 9).

**Figure 3.**
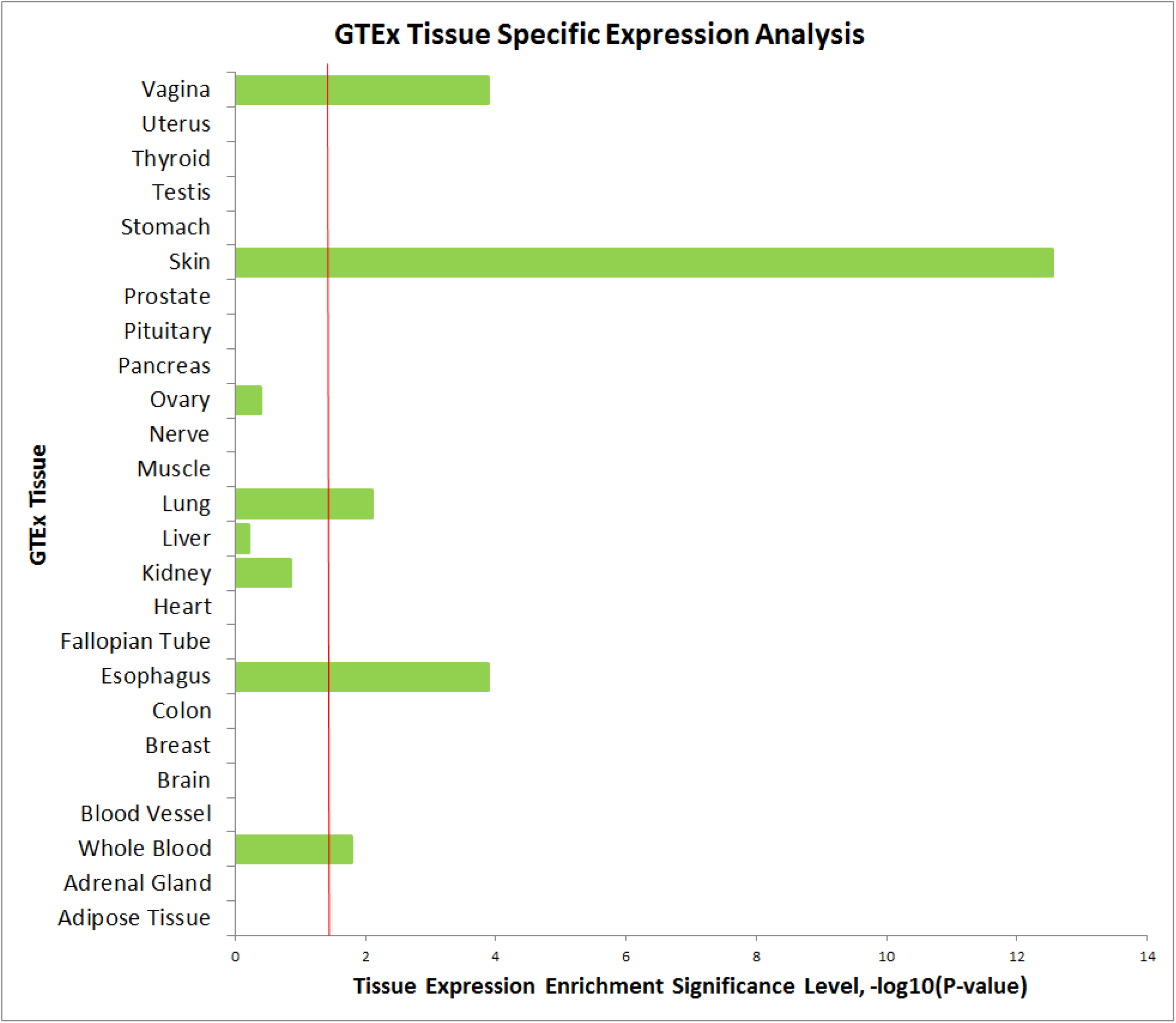
GTEx Tissue Specific Expression Analysis (TSEA). The horizontal axis illustrates tissue expression enrichment P-values after after Benjamin-Hochberg correction. The vertical axis illustrates 25 independent tissue types from GTEx.

In our replication analysis, we have confirmed a significant level of genetic correlation between asthma and eczema by using two independent GWAS data: the GABRIEL study^4^ and EAGLE study^27^. LD score regression results showed there was a strong positive genetic correlation between asthma (GABRIEL) and eczema (EAGLE) (*r_g_* = 0.46, *P* = 0.0044) (Table 1). We have then used CPMA to investigate the shared genes between asthma and one allergic disease, eczema. Thirteen loci demonstrated genome-wide significance. Of these, genes IL6R, IL1RL1, IL13, IL4, ACSL6, TSLP, IL33, C11orf30 and GSDMB were replicated comparing with the UK Biobank results (Supplementary Table 4). SMAD3 was found as a suggestive gene (rs744910, *P*_meta_ = 1.46×10^−7^) even though it does not reach to the genome-wide significance (Supplementary Table 5). While the phenotypes in our replication phase are not exactly same as UK Biobank’s, these results underscore the emerging overlap in genetic bases of asthma and allergy.

GO analysis highlighted several biological processes significantly enriched of the shared genes between asthma and allergic diseases, including epithelium development, keratinization, skin development and T helper 1 type immune response (Fig.4 and Supplementary Table 6). Also KEGG pathway analysis found the shared genes were significantly enriched in immune related pathway (Supplementary Table 7).

**Figure 4.**
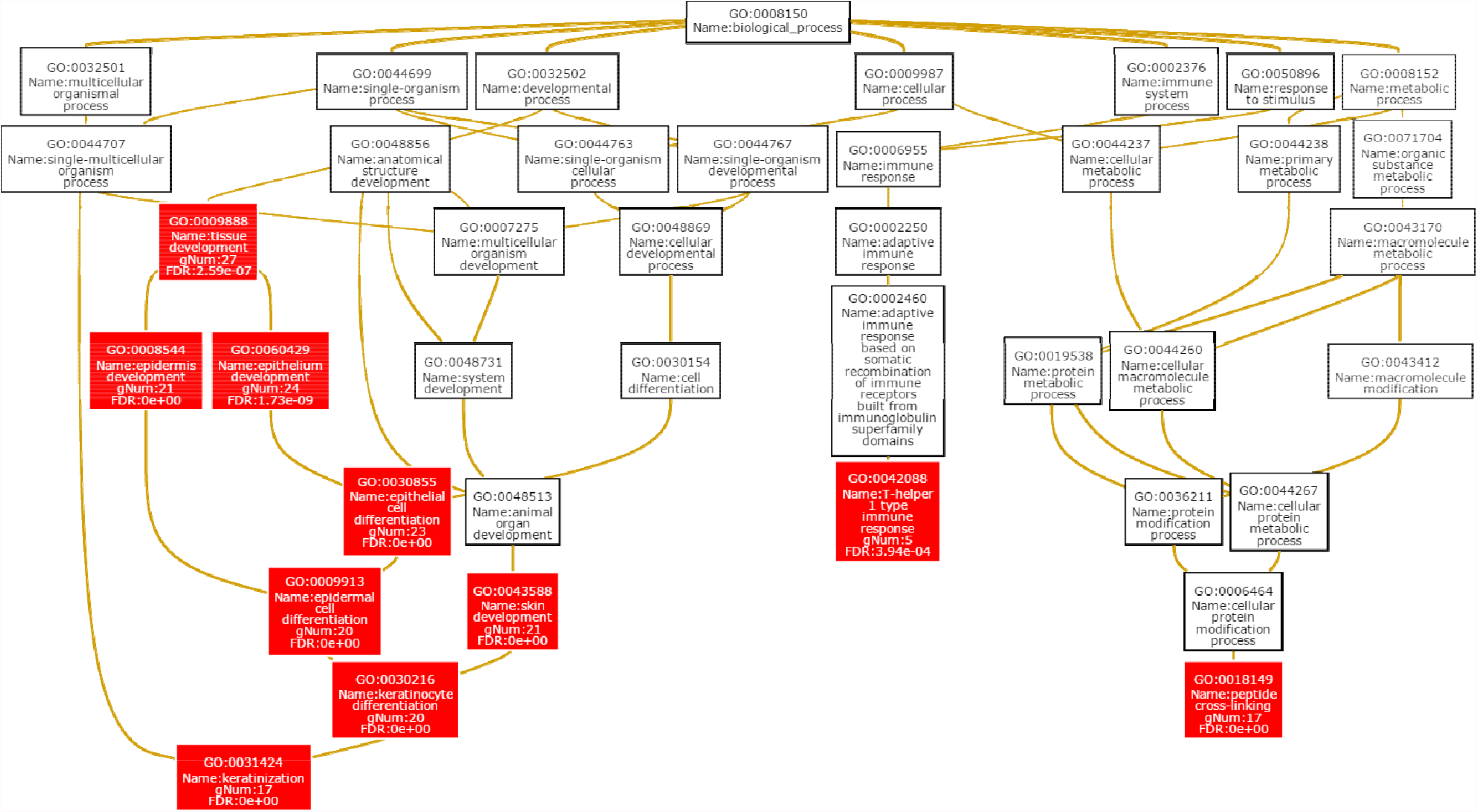
Gene Ontology (GO) biological process enrichment analysis. Red box indicates significantly regulated biological process (after false discovery rate).

## Discussion

In this study, we have identified strong genetic correlation between asthma and allergic diseases (hay fever/allergic rhinitis or eczema) and shared genetic loci between them. Further we have replicated our finding in two independent GWAS studies^4,27^. We have also found shared genetic loci between asthma and allergic diseases were significantly enriched in several tissues, including skin, esophageal, vagina, lung and whole blood tissue expression.

A total of 7 new loci were identified for asthma and 11 for allergic diseases. Of these, *D2HGDH* and *GAL3ST2* are associated with both asthma and allergic diseases, and have significant single tissue expression quantitative trait loci (eQTL) including lung, esophagus mucosa, skin (Supplementary Table 8 and 9). D2HGDH encodes D-2hydroxyglutarate dehydrogenase enzyme regulate alpha-ketoglutarate^52^ and can regulate alpha-ketoglutarate levels^59^, which also regulate Th1 cell and regulatory T cell^60^. Also Lian et al showed down-regulation of isocitrate dehydrogenase 2 (IDH2) is likely one of the mechanisms underlying altered 5-position of cytosine expression within epidermal basal cell layer^61^. The defection of such layer can also progress to asthma by releasing cytokine into systemic circulation and sensitizing to the lung^62^. GAL3ST2 encodes tumor metastasis related Galactose-3-O-sulfotransferase 2 enzyme53 and is expressed in the human small airway epithelium^63^. We also found rs12413578, near HV745896, a new regulatory genes for human telomerase reverse transcriptase (hTERT) gene expression, to be associated with both traits. Telomerase can be activated by antigen, leading to allergen-specific immune responses^64^. In terms of asthma, we found rs6461503, near integrin subunit beta 8 (ITGB8) as a novel loci. This gene was previously reported involving in T-cell mediated inflammation and autoimmunity^65,66^.

To diminish heterogeneity of significant loci potentially attributable to overlap samples in UK Biobank, we performed sensitivity analysis of excluding allergic asthma subjects (overlap sample between asthma and allergic diseases). When analyzing the 7,908 asthma and 19,508 allergic diseases subjects with 76,768 control subjects, we still observed significant genetic correlation between two traits (*r_g_* = 0.42, P = 6.64×10^−7^). In meta-analysis, most of the loci still showed robust genome wide shared associations between both traits, such as HLA-DQ, IL1R1, C11orf30, SLC25A46, D2HGDH and many others (Supplementary Table 10 and Supplementary Fig. 7). Although the significance level was reduced, this was mainly due to the loss of sample size for 6,177 individuals. In our sensitivity analysis for effect estimate, which should not be biased by sample size, we observed highly similar results between including and excluding allergic asthma subjects (Supplementary Table 10 and Supplementary Fig. 8).

We found genes that are associated with IgE function can consist of another crucial pathway for atopic diseases, such as asthma and eczema. We have found C11orf30 on chromosome 11 (Supplementary Table 2-4) as another shared genes between asthma and allergic diseases. C11orf30 was reported to be associated with total serum IgE levels and increases susceptibility to poly-sensitization^49^. When individuals are exposed to allergen, IgE are released from the immune system and travel to local organs or tissues to release type 2 cytokines such as IL-4,IL-13 and other inflammatory mediators^67^. These cytokines can further cause allergic diseases and asthma, such as reddish/dry skin and coughing, wheezing or shortness of breath^68,69^.

Clinical and epidemiologic observations have shown that immune-mediated inflammatory and autoimmune diseases can be associated due to overlap of genes^12^. Indeed, our LDSC analysis showed high genome-wide genetic correlation between asthma and allergic diseases and identified shared genes between them. However, we did not find significant genetic correlation between asthma and three other immune related diseases, rheumatoid arthritis, Crohn’s disease and ulcerative colitis, suggesting that the patterns of genetic architecture among immune related traits can still be distinct. Asthma and allergic diseases both belong to IgE mediated hypersensitivity (Type I). While the other three traits are immune related, rheumatoid arthritis belongs to immune complex mediated (Type III) hypersensitivity, and Crohn’s disease and ulcerative colitis belong to delayed cell mediated (Type IV) hypersensitivity. Thus we found strong genetic correlation for the traits within each hypersensitivity category. Different types of hypersensitivity are characterized by different types of gene sets.

Though the association between respiratory diseases like asthma and skin tissue diseases may, at first, seem counterintuitive, exploring the link between the two apparently distinct phenotypes from a gene function perspective reveals a noticeable overlap in underlying genetics and molecular pathways in their etiologies. We have confirmed FLG gene on chromosome 1 (Supplementary Table 2-4) was associated with both allergic diseases and asthma. FLG is important for the formation of the stratum corneum and also for the hydration of this barrier layer^58^. People who have mutations in the FLG gene can be sensitive to external allergens and produce dry and flaky skin^58^. Meanwhile it can activate a strong allergic immune response through peripheral blood to many organs, such as lung, causing the inflammation in the lungs that results in shortness of breath^58^.

Our predominant finding in whole blood tissue enrichment indicated that these diseases were caused by some malfunction of the shared immune system. Genes such asCD247 and RUNX3 play important roles in T-helper cells, CD8^+^ T cells, NK and DCs ^70^ were previously reported as a risk variant for many immune-related inflammatory diseases, such as asthma, Crohn’s disease, ulcerative colitis and eczema^70-73^.

Our findings suggest that the shared pathway between asthma and allergic diseases might have significant functions in tissues like skin, esophagus, vagina, lung and whole blood. Skin, esophagus, vagina and lung tissues are all characterized by various type of epithelium or mucosa^74,75^, which share similarities in terms of functions and mechanisms. For example, the epithelial cells lining the airway, lungs and skin serve as the primary defense to inflammatory stimuli and antigens infiltrating these tissues through inducing cell signaling pathways in response to contact with the irritants. Indeed, a hallmark of asthma and allergic rhinitis is allergic sensitization by means of epithelial activation in the respiratory tract^76,77^. Also epithelial cells can transdifferentiate into motile mesenchymal cells, which can behave like stem cells and differentiate into multiple cell lineages and migrate to damage tissues to repair^78^. Thus, the enrichment of asthma and allergy genes expressed in epithelial components is not specific to a certain organ or tissue; rather, it seems to be generalizable across epithelium types. Also whole blood is responsible for production of anti-inflammatory cytokines and circulating them in the whole body^79^.

We also acknowledge the limitations of our study. First, more allergic diseases will be needed to investigate if genes are specific to certain types of allergy shared with asthma, such as food allergy. Also, our allergic symptom phenotype contained both allergic rhinitis/hay fever and eczema. The allergy genes we found may not be associated with a specific type of allergy.

However, since allergic rhinitis and eczema are both type I hypersensitivity and characterized by epithelial cells, they should share highly similar physiology^80,81^. Finally, our UK Biobank cohort traits are not independent. There are shared cases and complete shared controls between asthma and allergic diseases. However, for all of our analysis, we have applied methods that can adjust potential effect due to overlapping samples; and we have also applied the sensitivity analysis and two independent GWAS data to approve the robustness of shared loci. A surprising finding in replication analysis was we did not observe a significant association in the HLA region. This may be due to different GWAS platforms in the GABRIEL and EAGLE studies, or that 1000 genome imputation was conducted for GABRIEL GWAS summary statistics. This process provided fewer total SNPs that are common for both traits than in UK Biobank, which could be another reason that only part of the shared genes were found in this analysis.

These shared genetic components can provide opportunity to investigate asthma and allergy comorbidity on molecular level. Drugs targeting shared genetic loci might benefit treatment of both diseases. Our findings can also inspire patients and doctors to be more cautious about comorbidity of asthma and allergic diseases in the clinical practice.

## Conclusion

Understanding the genetic overlap between asthma and allergic diseases can be beneficial to treat disease outcome more efficiently. Our GWAS study has highlighted shared genetic pathway in immune and inflammatory systems and epithelium tissues by asthma and allergic diseases. This work reinforces the idea that asthma and allergy implicate shared common biological processes and open new avenues of treatment for them.

URLs. 1000 Genomes Project, http://www.1000genomes.org/; BEAGLE, http://faculty.washington.edu/browning/beagle/beagle.html; UK Biobank, http://biobank.ctsu.ox.ac.uk/; LiftOver, http://genome.sph.umich.edu/wiki/LiftOver; ImpG, https://github.com/huwenboshi/ImpG; LDSC, https://github.com/bulik/ldsc; GTEx, http://www.gtexportal.org; TSEA, http://genetics.wustl.edu/jdlab/tsea/; WebGestalt, http://www.webgestalt.org/option.php; PLINK, https://www.cog-genomics.org/plink2; GenABEL, http://www.genabel.org/packages/GenABEL; R, https://www.r-project.org/; GWAS catalog, https://www.ebi.ac.uk/gwas; ASSET, https://bioconductor.org/packages/release/bioc/html/ASSET.html; CPMA, http://coruscant.itmat.upenn.edu/software.html; GCTA, http://cnsgenomics.com/software/gcta; GABRIEL consortium, https://www.cng.fr/gabriel/index.html; EAGLE consortium, http://data.bris.ac.uk/data;

## Authors’ contributions

ZZ, LL, PHL, QL, DCC designed the study.

ZZ, LL, MC, WC, PRL performed the statistical analysis. ZZ, LL, WC, MC, QL wrote the manuscript.

All authors helped interpret the data, reviewed and edited the final paper, and approved the submission.

ZZ and LL had full access to all the data in the study and takes responsibility for the integrity of the data and the accuracy of the data analysis.

## Competing interests

The authors declare no competing interests.

## Acknowledgments

This research has been conducted using the UK Biobank Resource under application number 16549. We would like to thank the participants and researchers from the UK Biobank who significantly contributed or collected data. We thank to GABRIEL consortium and EAGLE consortium for providing GWAS summary statistic data. We also thank to Dr Vernerri Anttila and Dr Steven Gazal for their statistical advice. We thank Elizabeth A. Loehrer for editorial assistance.

